## Supplementary Material for "Programmable and portable CRISPR-Cas transcriptional activation in bacteria"

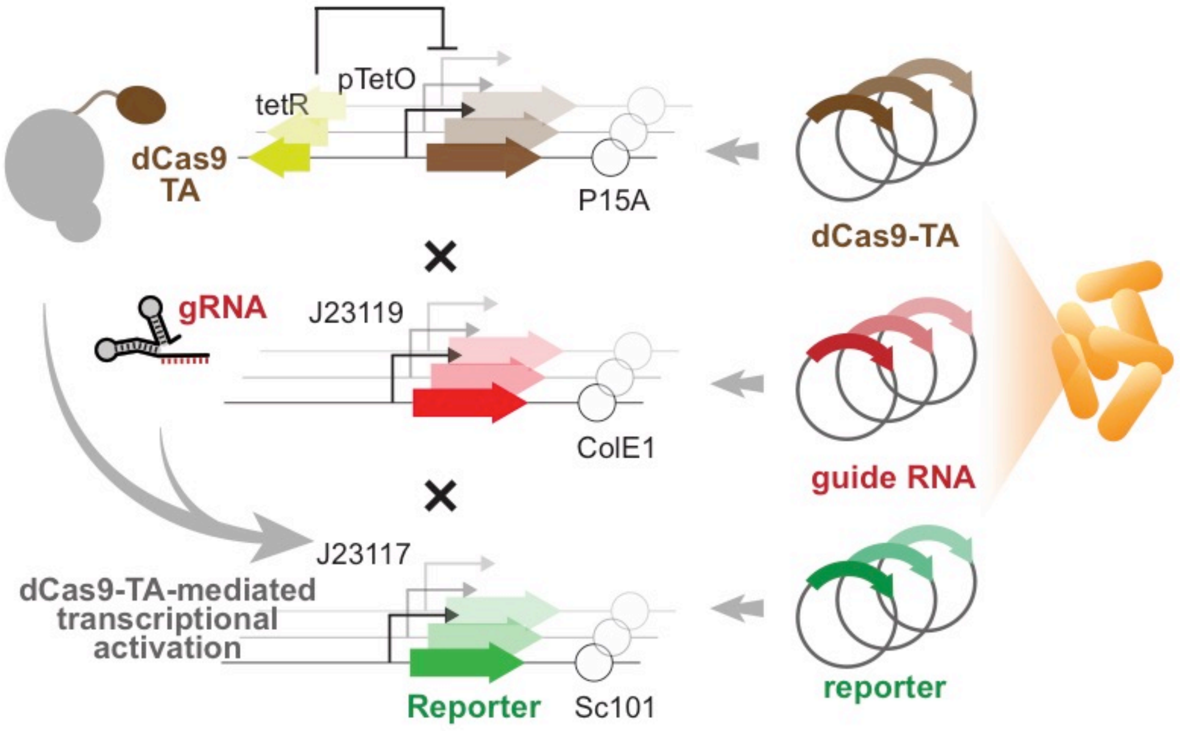


**Figure S1. Diagram of CasTA platform** The design of separating 3 key components of CRISPRa, dCas9-TA, gRNA, and reporter, into 3 compatible plasmids that could function in the same cell.

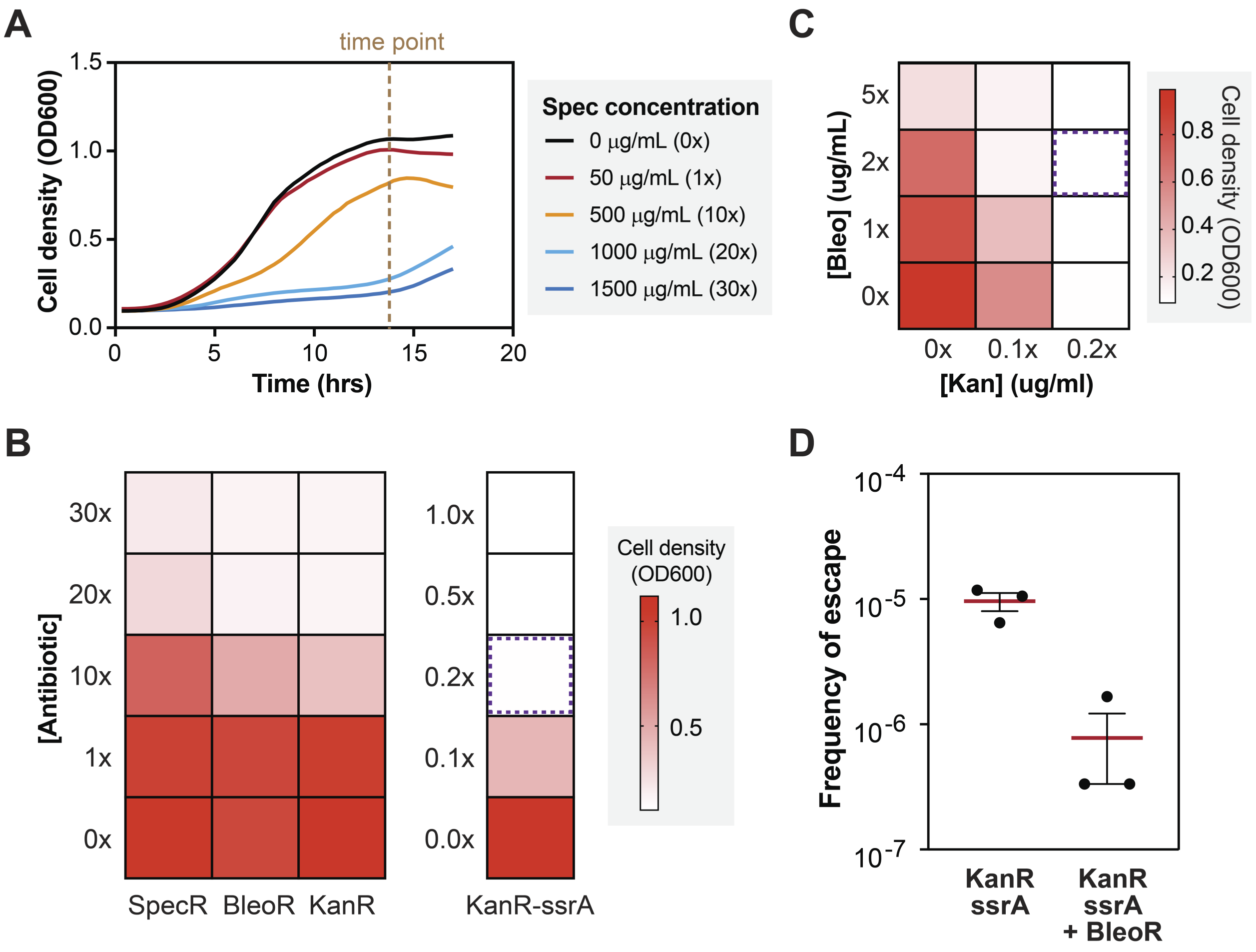


**Figure S2. Optimization of selection stringency for CasTA selection platform. (A)** Growth curves of *E. coli* containing pHH34 on LB media supplemented with different spectinomycin concentrations. Dotted line indicates growth phase when cell density was measured in other panels. **(B)** Selection stringency of different antibiotics using corresponding resistance genes as selection reporters (pHH34-37). KanR-ssrA: Kan resistance gene (KanR) with degradation tag (AANDENYALAA). Heat map correspond to cell density after 14 hrs. Purple dotted outline correspond to the antibiotic concentration used for sufficiently stringent selection. For SpecR, 1x Spectinomycin = 50 μg/ml. For BleoR, 1x Bleocin = 5 μg/ml. For KanR, 1x Kanamycin = 50 μg/ml. **(C)** Selection stringency of KanR-ssrA (x-axis) and BleoR (y-axis) dual reporter with double antibiotic selection of Kanamycin (Kan) and Bleocin (Bleo). Purple dotted outline correspond to the antibiotic concentration used for sufficiently stringent selection. **(D)** Escape rates of using KanR-ssrA alone or KanR-ssrA and BleoR as selection reporters. Data are 3 biological replicates in each experiment. Errorbars are S.E.M.

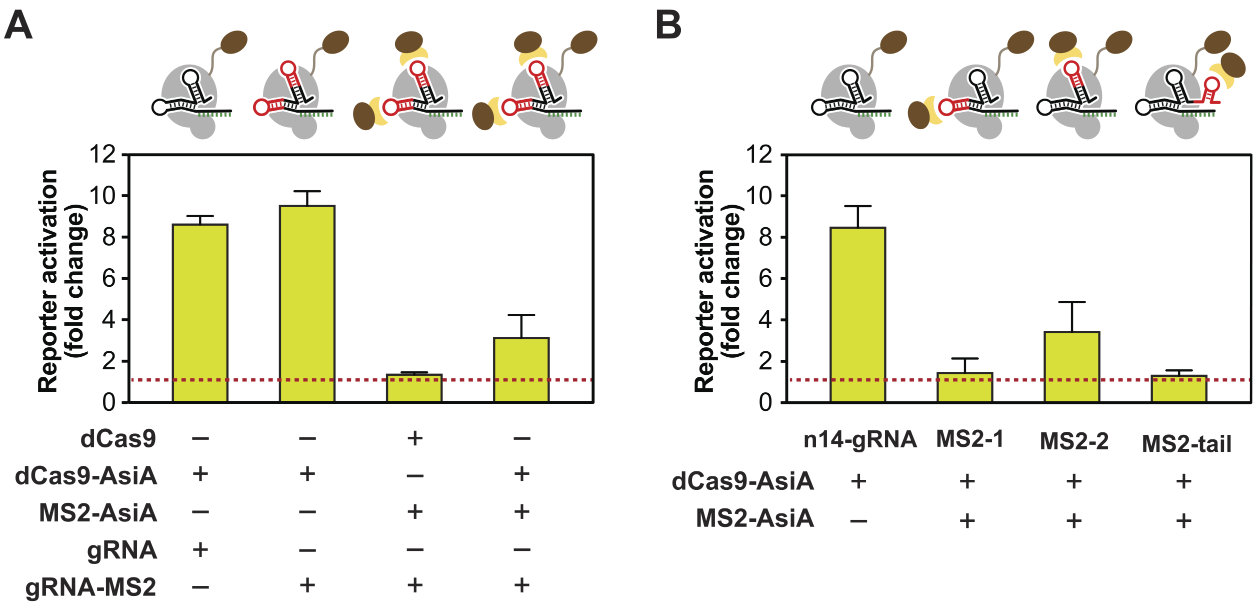


**Figure S3. Evaluation of different dCas9 transcription activator fusion strategies. (A)** dCas9 SAM system with modified gRNA and MS2-AsiA does not enhance CRISPRa activity. dCas9 tether AsiA is required for facilitating gene activation. **(B)** Examination of different gRNA designs for improving CRISPRa. n14-gRNA represents design with only 14 nucleotides of the N20 seed sequence; MS2-1: incorporating MS2 hairpin structure in the first loop of the wild-type gRNA structure; MS2-2: incorporating MS2 in the second loop of the wild-type gRNA structure; MS2-tail: MS2 was fused at the 3' end of the gRNA structure. Bars are mean of 3-5 biological replicates with errorbars showing as S.E.M.

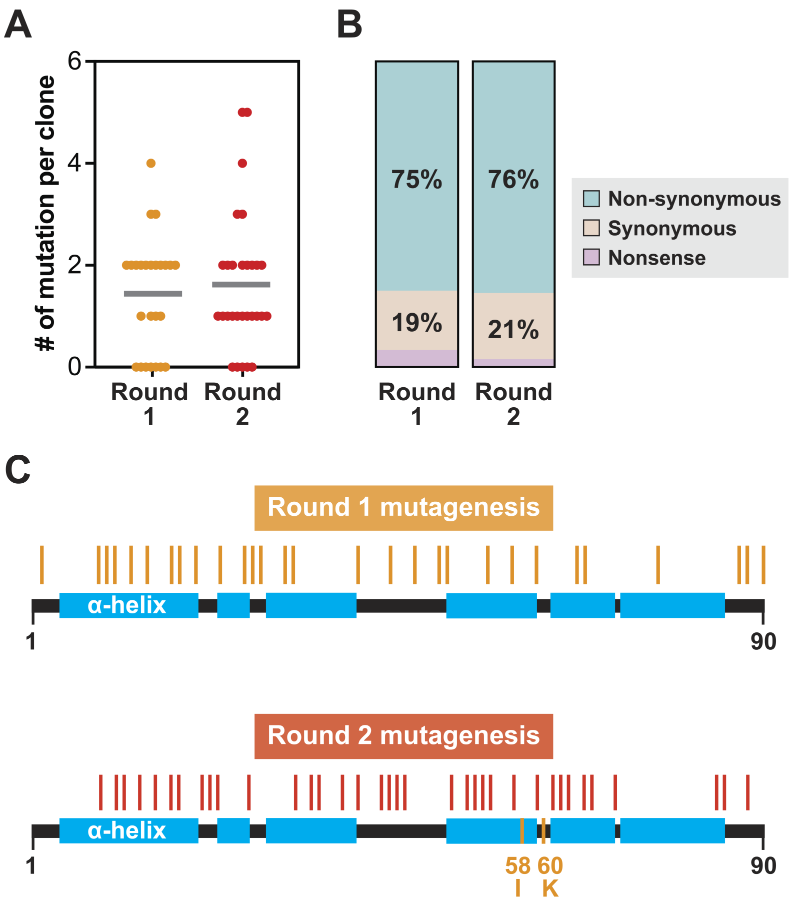


**Figure S4. Sequence profiling of AsiA variant libraries after PCR mutagenesis.** Sanger sequencing of AsiA variants after 2 rounds of mutagenesis **(A)** the number of mutations per variant, **(B)** types of mutations in the protein sequences, and **(C)** mutated positions along the protein secondary structure of AsiA (indicated by colored ticks). Profiles are generated based on at least 25 randomly selected variants from each round of mutagenesis.

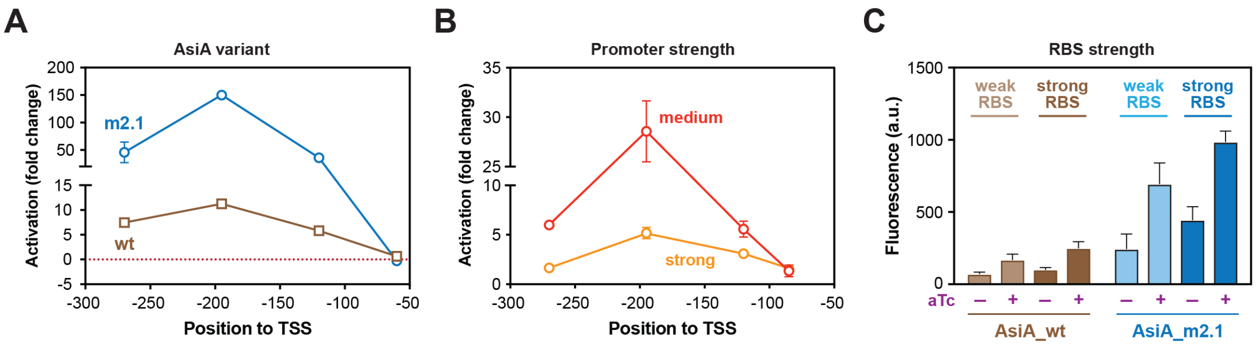


**Figure S5. Characterization of dCas9-AsiA mediated CRISPRa. (A)** Transcriptional activation of a weak promoter (J23117) of a GFP reporter using different gRNAs with dCas9-AsiA wild-type (wt) or mutant (m2.1). dCas9-AsiA_m2.1 (blue circles) has similar optimal gRNA targeting distance (~-200bp from TSS) as dCas9-AsiA_wt (brown squares). **(B)** Transcriptional activation using dCas9-AsiA_m2.1 with different gRNAs against a medium basal strength promoter (J23116; red circles) or strong basal strength promoter (J23110; orange circles). Induction range is found to be higher for the medium promoter than the strong promoter due to saturating absolute induction level for both promoters. **(C)** Increasing ribosomal binding site (RBS) strength (**Table S9**) with and without transcriptional induction (+/- aTc) of dCas9-AsiA wild-type (wt) or mutant (m2.1) generally increased fluorescence signal of reporter gene. Weak RBS (BBa_B0033), strong RBS (BBa_B0034). Mean from three biological replicates are plotted with errorbars as +/- S.E.M.

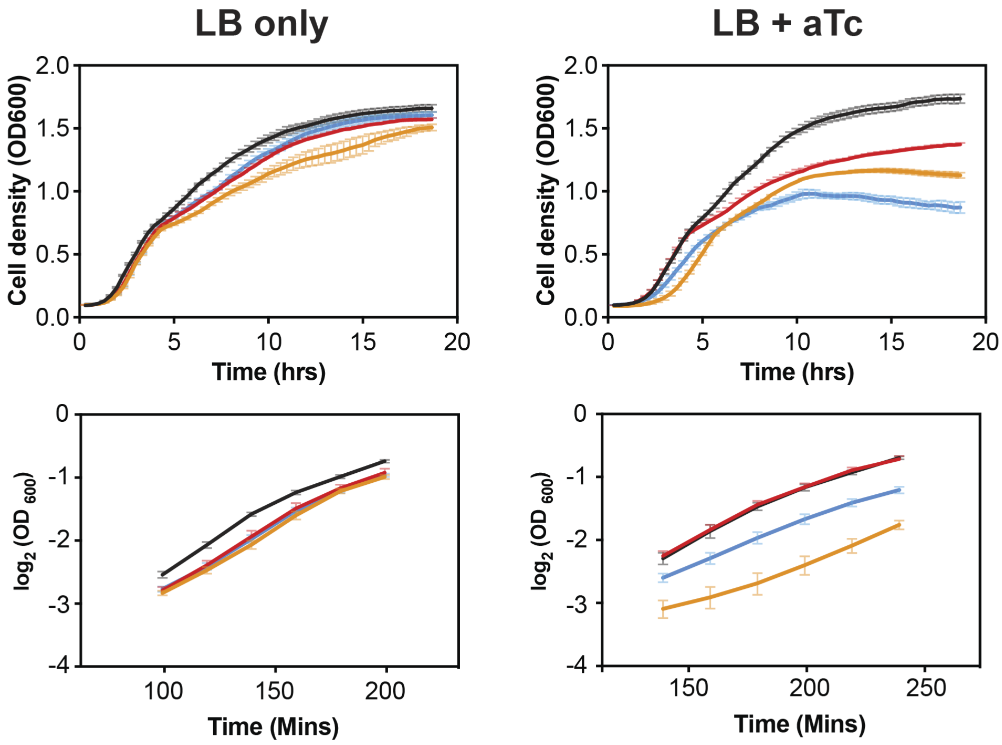


| | doubling time (mins) | |
|---|---|---|
| | LB only | LB + aTc |
| BW25113 (WT) | 57-68 | 62-77 |
| dCas9 | 50-64 | 63-81 |
| dCas9-AsiA | 51-60 | 62-77 |
| dCas9-AsiA_m2.1 | 53-62 | 65-85 |

**Figure S6. Growth of cells expressing dCas9-AsiA.** Cells carrying different dCas9-AsiA plasmids were grown in rich media with (LB+aTc) or without (LB only) dCas9 overexpression. Growth curve and doubling times in the exponential growth phase are shown. Data are three biological replicates with errorbars as +/- S.E.M.

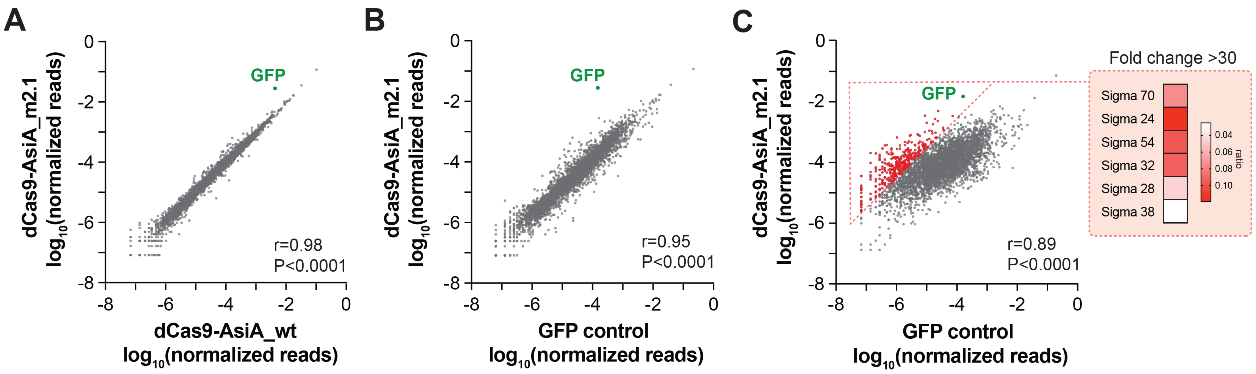


**Figure S7. Specificity of gene activation using dCas9-AsiA_m2.1. (A)** Transcriptomic profile of cells expressing dCas9-AsiA_wt using an optimal gRNA (gRNA-H4) targeting a GFP reporter gene on pWJ89 (x-axis) versus cells expressing dCas9-AsiA_m2.1 and the same gRNA (y-axis). **(B)** Transcriptomic profile of parental GFP control (pWJ89) cells (x-axis) versus cells expressing dCas9-AsiA_m2.1 and gRNA-H4 (y-axis). **(C)** Transcriptomic profile of parental GFP control cells (x-axis) versus with cells overexpessing dCas9-AsiA_m2.1 and gRNA-H4 (y-axis). Genes with more than 30 fold up-regulation under dCas9-AsiA_m2.1 over-expression are highlighted in red and grouped by their annotated sigma factors. Heatmap on the right indicates the ratios of highly activated (fold change >30) promoters within each group of promoters mediated by different sigma factors.

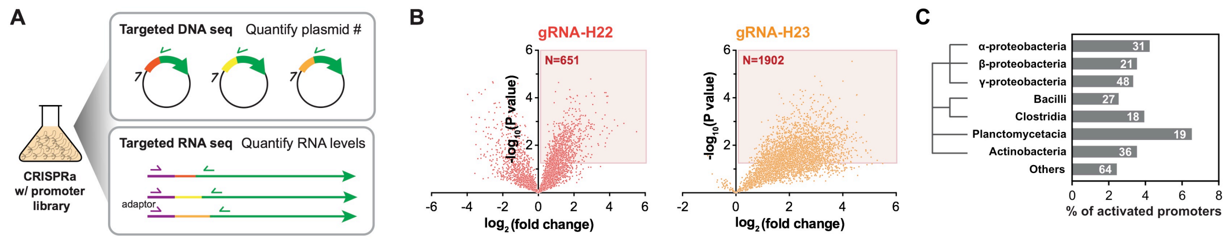


**Figure S8. Bacterial CRISPRa screen to identify new orthogonal inducible promoters. (A)** Using CRISPRa on a metagenomic promoter library (RS7003) to mine CasTA-inducible promoters using targeted DNAseq and targeted RNAseq. **(B)** Volcano plots of CasTA-mediated activation using two different gRNAs (gRNA-H22 and gRNA-H23) of the same promoter library, with each point in the plot corresponding to a unique promoter. Significantly activated promoters ($p<0.05$) are highlighted with the red rectangle, and the numbers of activated promoters are indicated. Data are calculated from 4 biological experiments. **(C)** The percentage of highly activated promoters (fold change >10) among all promoters of each bacterial genius. Numbers in the bars indicate the actual numbers of highly activated promoters. Dendrogram represents the phylogenic distance between each group.
